# Antibody Humanization via Protein Language Model and Neighbor Retrieval

**DOI:** 10.1101/2023.09.04.556278

**Authors:** Honggang Zou, Rongqing Yuan, Boqiao Lai, Yang Dou, Li Wei, Jinbo Xu

## Abstract

Antibody (Ab), also known as immunoglobulin (Ig), is an essential macromolecule involved in human immune response and plays an increasingly vital role in drug discovery. However, the development of antibody drugs heavily relies on humanization of murine antibodies, which often necessitates multiple rounds of sequence optimizations through laborious experimental processes. In recent years, the remarkable capabilities of machine learning have revolutionized the field of natural sciences and have also demonstrated promising applications in the field of antibody humanization. Here, we present Protein-LAnguage-model-knN (PLAN), a machine learning model leveraging protein language model and information retrieval for improving humanization of antibodies. Further, we propose *D*_*E*_, a computed value shows a positive correlation with antigen-binding affinity. Our *in silico* experimental results demonstrate that 1) the PLAN-humanized sequences’ average humanness score reaches 0.592, improving over the best existing method by 44.7%; 2) a 63% overlap between the PLAN-proposed mutations and the mutations validated through wet lab experiments, which is 16.7% higher than the best existing result; 3) comparable antigen-binding affinity after *D*_*E*_ guided back mutation.

## Introduction

Antibody (Ab), also known as immunoglobulin (Ig), contains two identical heavy chains and two identical light chains connected by disulfide bonds. Each heavy and light chain contains three hypervariable regions, giving rise to three loops of β-strands. Three hypervariable regions, which are referred to as Complementarity Determining Regions (CDRs [6]), from each of the heavy and light chain pairs together form the paratope of the antibody. Antibody numbering system such as Kabat, IMGT, and Chothia [15][16] are used to accurately identify and compare CDRs from a given antibody heavy or light chain sequence. Computational tool such as ANARCI [17] aligns a given sequence to a database of Hidden Markov Models that describe the germline sequences of antibody and TCR domain types, thereby assigning the corresponding CDR numberings.

Due to the capability of antibody to specifically bind to antigen with high affinity, an antibody can tag a microbe for phagocytosis or can neutralize it directly. Thus, antibody plays an important role in human immune response and has the potential to be developed as drug for the therapy and diagnosis of different human diseases [32]. Murine antibody has drawn much attention for clinical use due to its availability, low cost, and quick production time [20]. Nevertheless, the introduction of murine antibodies into patients may trigger a human anti-antibody response (AAR). This phenomenon can significantly diminish the intended therapeutic impact while potentially inducing adverse systemic inflammatory reactions in patients. This occurrence, often referred to as antibody immunogenicity, underscores the importance of addressing such challenges in antibody-based therapeutics development. Various methods have been developed to humanize parental murine antibodies, by weakening its immunogenicity while preserving its essential biophysical properties, such as antigen-binding affinity through sequence optimization [1].

Antibody humanization methods can be mainly divided into two categories (Figure 1): Grafting and Mutating. Grafting, including CDR grafting [4], SDR grafting [5], and energy-based grafting method [26], is to graft functional part of murine antibody onto a human antibody scaffold, thereby reducing antibody’s immunogenicity while retaining its antigen-binding affinity. Mutating can be further divided into humanness scoring-based method and protein language model-based method, where the former introduces mutation by scoring antibody’s immunogenicity, while the latter introduces mutation by modeling antibody using protein language models.

**Figure 1.**
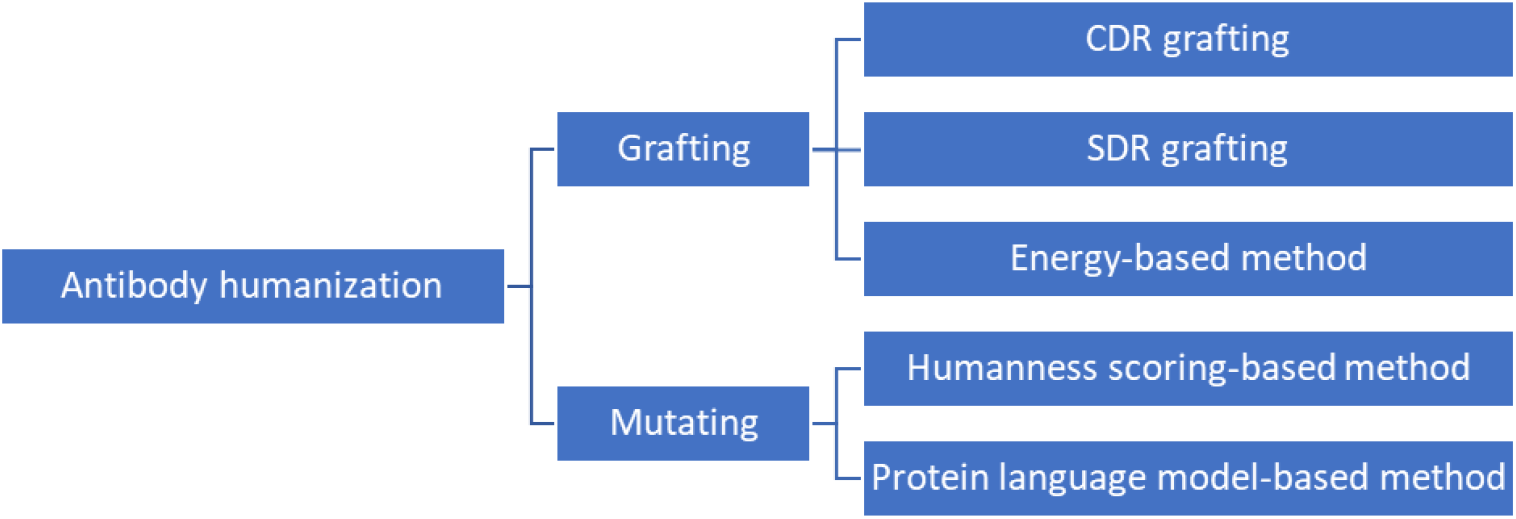
Antibody humanization methods classification.

The most widely used approach to humanizing murine antibody is CDR grafting [4]. CDR grafting is to graft the CDRs of the parental murine antibody onto a carefully selected human antibody. However, retention of the CDRs is often not sufficient to retain antigen-binding affinity [1], since non-CDRs residues may contribute to the structural conformation of CDRs. *In vitro* experiments are then carried out to locate and back mutate those key non-CDRs residues. In general, CDR grafting has the advantages of easy implementation, but often laborious and time consuming.

Several computational metrics have been developed to distinguish murine and human antibodies by computing a humanness score for antibodies. For example, human string content (HSC) score was calculated by determining the proportion of peptide strings within an antibody that could also found within a set of human antibodies [7][8]. Inspired by HSC score, OASis calculates the fraction of peptides within a single antibody that pass a user-defined prevalence threshold [10]. On the other hand, predictive models can also be used to score antibody humanness. For example, deep learning model based on bi-directional long short-term memory (LSTM) architecture was trained to classify murine and human antibody [22]. These metrics could be utilized in a humanness scoring-based humanization method to substitute for time-consuming and trial-and-error *in vitro* experiments to reduce immunogenicity [7][8][9][10][22]. *In silico* humanization protocol such as Hu-mAb [9] was implemented by introducing mutations to sequence and measuring the humanness score iteratively until the humanness score surpasses a pre-defined threshold. However, they do not consider the antigen-binding affinity. One feasible mitigation is to minimize the number of mutations made to the sequence to limit the unpredictable impact on affinity, but the weakness remains.

Natural language processing (NLP) models have given new momentum to the computational antibody design field [21]. It is intuitive to regard the amino acid sequence as sentence and amino acid as token, so that a large number of NLP models can be applied to protein modeling. Thanks to the next generation sequencing (NGS) and other advanced biological technologies, a very large-scale experimental data could be fed to NLP models [14]. Using a self-supervised masked language modeling (MLM) task, several pre-trained protein language models (PLM) have been developed [10][11][12][13][31]. The representations, also called embeddings, of proteins generated by PLM can then be utilized as inputs for downstream tasks, including humanization of antibodies. For example, Sapiens [10], a PLM pre-trained on OAS [23], can recognize and repair non-human residues, thereby humanizing murine antibodies. Capitalizing on the information-rich protein embeddings extracted from PLM to effectively humanize murine antibody necessitates access to a substantial dataset of murine antibody-humanized antibody pairs. However, the availability of such datasets remains limited. To overcome this limitation, Sapiens [10] was trained on human antibody database and performed masked language modeling (MLM) on murine sequences, enabling Sapiens to humanize the masked amino acid, hence avoiding supervised training. Different from Sapiens, we leverage distance-based neighbor retrieval approach, such as k-nearest neighbor search, in the embedding space to identify potential humanization candidates.

K-nearest neighbor model is a conventional machine learning algorithm with great interpretability, based on the principle that adjacent points in the representation space have similar properties. Previous study [24] extend a pre-trained language model by linearly interpolating it with a kNN model. Subsequent study [25] show that this idea is beneficial for open-domain question answering (QA). In this study, we demonstrate that this BERT-kNN paradigm can also be applied to the humanization of antibodies. In this study, we developed a novel machine learning method called PLAN (Protein-LAnguage-model-knN) for antibody humanization. Different from previous work where language models [33] (or generative models [34]) were used to output probabilities of mutated amino acids (or protein sequences) to guide protein optimization, PLAN employs protein language model to extract the feature representation for each amino acid in the input murine sequence, and then runs kNN search in representation’s space to search for amino acid mutations from the human antibody repertoire, thereby humanizing the input sequence. *In silico* experiments demonstrate that 1) the PLAN-humanized sequences’ average humanness score reaches 0.592, improving over the best existing method by 44.7%; 2) a 63% overlap between the PLAN-proposed mutations and the mutations validated through wet lab experiments, which is 16.7% higher than the best existing result; 3) comparable antigen-binding affinity after *D*_*E*_ guided back mutation. Additionally, *in vitro* and *in silico* experiments confirms that our method can identify residues that have a critical impact on affinity, thereby providing valuable guidance for back mutation.

## Results

A list of 25 experimentally humanized therapeutic sequences and their precursors was created by [9]. These 25 antibodies can perform as a validation set for our humanization protocol. Analogous to [9][10], we humanize these 25 antibodies with our protocol and measure the number of mutations that were both suggested by our protocol and made experimentally. Since our method sorts all 20 amino acids at each position, we can also calculate top 1, top 2, and top 3 accuracy on residue level, regarding the precursor as input, the humanized candidates generated by our method as output, and the experimentally humanized sequence as ground truth. We utilize OASis [10] and IgBLAST [27] to evaluate the humanness score and Flex ddg (default settings) [30] to evaluate the antigen-binding affinity of antibody, and compare between PLAN, Hu-mAb [9], and Sapiens [10]. It is noted that in the following text, Sapiens (1 iteration) performed one round of Sapiens optimization, while Sapiens (3 iteration) performed three rounds (take the output of the previous round as input for each round).

The pipeline and evaluation method for PLAN are demonstrated in Figure 2.

**Figure 2.**
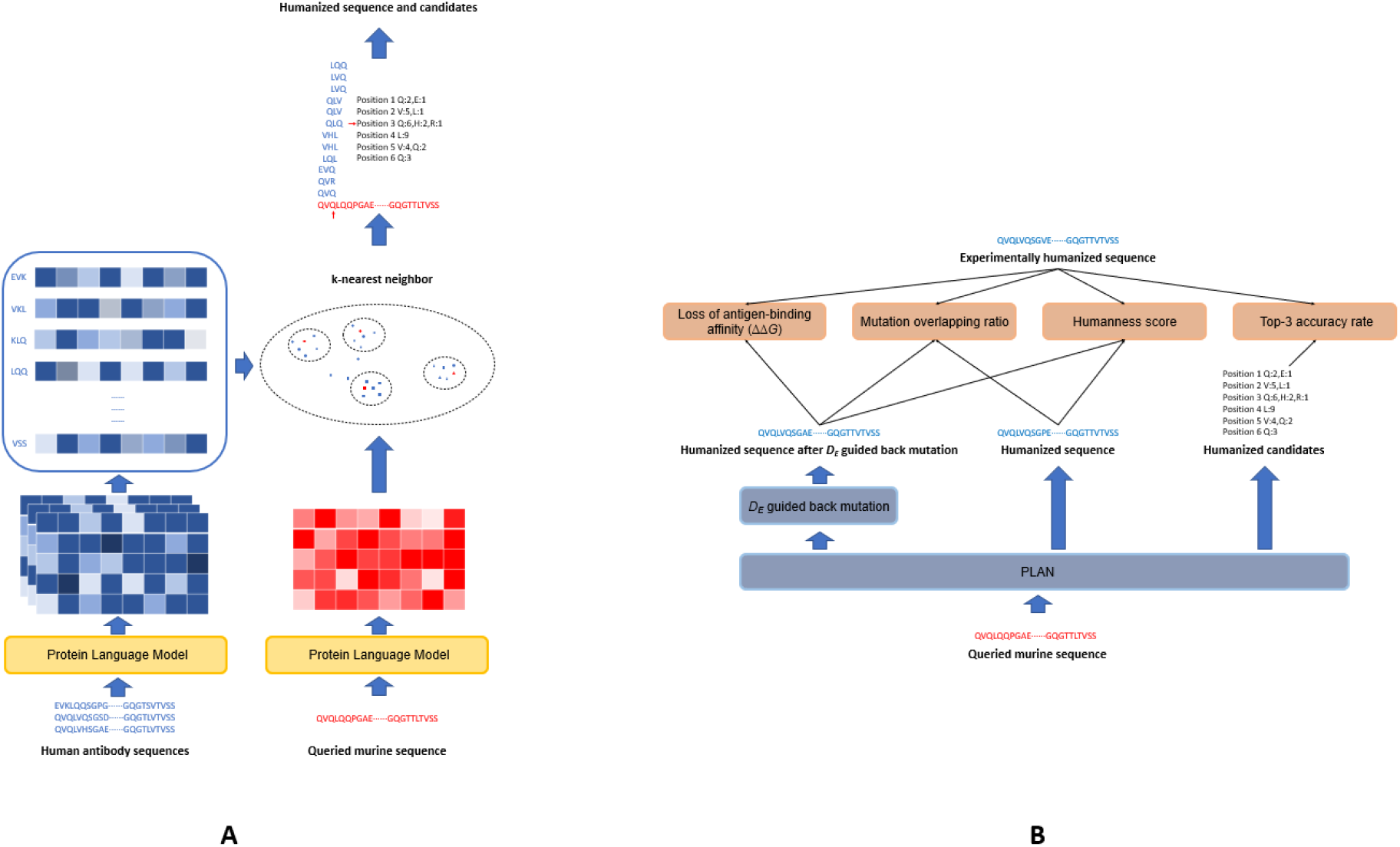
Pipeline and evaluation method for PLAN. (A) PLAN first collects numerous human antibody sequences and extracts their representations using protein language model (i.e., ESM2). The kNN pool is then established regarding each *K-mer* in collected sequences as key and corresponding embedding as value. For each embedding of *K-mer* in input murine antibody sequence, run neighbor retrieval (i.e., kNN algorithm) and obtain *k K-mers* from the kNN pool. Count the occurrence of all searched *K-mers* on residue level, and then vote for each position, selecting the amino acid with the highest count as the result of humanization. (B) Loss of antigen-binding affinity (ΔΔ*G*), mutation overlapping ratio, humanness score, and top-3 accuracy rate were measured to evaluate the effectiveness of PLAN.

### 1. PLAN successfully predicts mutations made by experimentally humanized sequences

We humanize the 25 antibodies with our protocol and measure the number of mutations that were both suggested by our protocol and made experimentally. For individual result, see the SI.1. For heavy chain, our method successfully predicts 380 out of 628 mutations made experimentally, accounting for 61%. For comparison, Hu-mAb [9], Sapiens [10] (1 iteration), and Sapiens (3 iterations) predicts 247, 238, and 275 out of 628 mutations made experimentally, accounting for 39%, 38%, and 44%, respectively. For light chain, our method successfully predicts 307 out of 469 mutations made experimentally, accounting for 66%. For comparison, Hu-mAb, Sapiens (1 iteration), and Sapiens (3 iterations) predicts 206, 264, and 312 out of 469 mutations made experimentally, accounting for 44%, 56%, and 67%, respectively. Overall, our method successfully predicts 687 out of 1097 mutations made experimentally, accounting for 63%. For comparison, Hu-mAb, Sapiens (1 iteration), and Sapiens (3 iterations) predicts 453, 502, and 587 out of 1097 mutations made experimentally, accounting for 41%, 46%, and 54%, respectively. The above results are also listed in Table 1 and demonstrated in SI.2.

**Table 1.** Result for mutation overlapping ratio between experimentally humanized sequences and PLAN-, Hu-mAb-, and Sapiens-humanized sequences among 25 antibodies.

| Method | Heavy chain | Light chain | Overall |
| --- | --- | --- | --- |
| PLAN | 61% | 66% | 63% |
| Hu-mAb | 39% | 44% | 41% |
| Sapiens (1 iteration) | 38% | 56% | 46% |
| Sapiens (3 iteration) | 44% | 67% | 54% |

It is worth noting that for Refanezumab, a monoclonal antibody designed for the recovery of motor function after stroke, PLAN-humanized light chain sequence is identical to the experimentally humanized light chain sequence (Figure 3).

**Figure 3.**
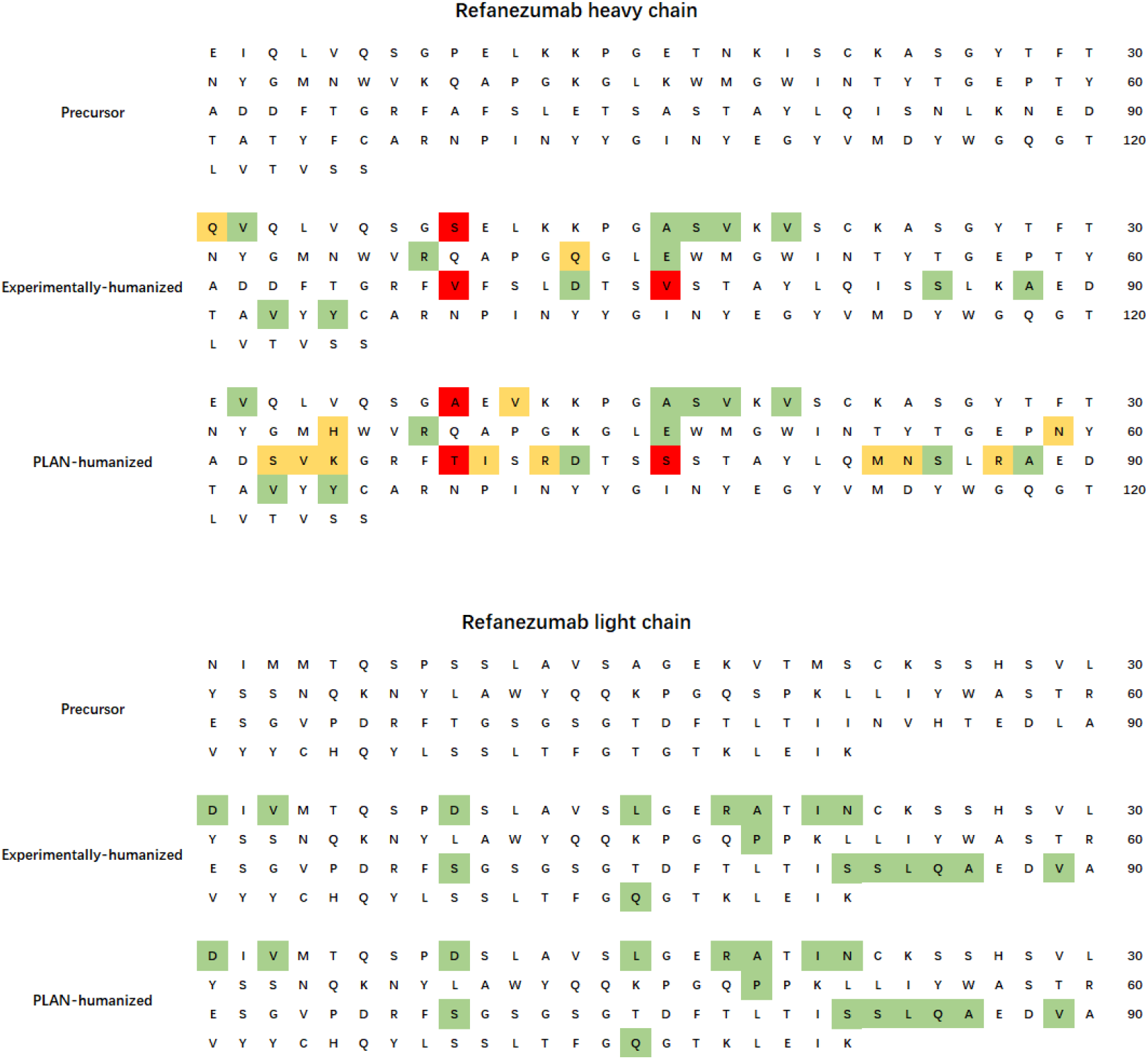
Mutation comparison chart for experimentally-humanized sequence and PLAN-humanized sequence of Refanezumab. The position stained green indicates identical mutation of experimentally-humanized sequence and PLAN-humanized sequence; The position stained red indicates different mutation of experimentally-humanized sequence and PLAN-humanized sequence; The position stained yellow in experimentally-humanized sequence (or PLAN-humanized sequence) indicates that the corresponding amino acid mutated in experimentally-humanized sequence (or PLAN-humanized sequence), but not in PLAN-humanized sequence (or experimentally-humanized sequence).

### 2 *In vitro* validation of Δ*D*_*E*_ guided back mutation

For a more detailed analysis, we classify each position into five categories (to exemplify this, SI.3 visualizes the results of classifying each position in an example):

i. (*Exp* = *Med*) & (*Exp* ≠ *Pre*).
ii. (*Exp* = *Med*) & (*Exp* = *Pre*).
iii. (*Exp* ≠ *Med*) & (*Exp* = *Pre*).
iv. (*Exp* ≠ *Med*) & (*Exp* ≠ *Pre*) & (*Med* ≠ *Pre*).
v. (*Exp* ≠ *Med*) & (*Exp* ≠ *Pre*) & (*Med* = *Pre*).

Where *Pre, Exp*, and *Med* denotes the amino acid at the corresponding position of precursor, experimentally humanized sequence, and sequence humanized by corresponding method, respectively.

The classification result for PLAN, Hu-mAb, and Sapiens among 25 antibodies is shown in SI.4.a (heavy chain) and SI.4.b (light chain). Since these methods do not introduce mutations in CDRs, we only consider non-CDR residues.

Although PLAN successfully predicts over three-fifths of the mutations made by experimentally humanized sequences, PLAN introduces approximately 12 extra mutations (iii + iv) per heavy chain and 7 extra mutations per light chain. This increases the risk of a decrease in antigen-binding affinity. This section discusses how to design back mutation to correct these mutations.

Intuitively, the representations of antibodies extracted from protein language model are related to their functions, so the difference of two sequence representations is related to their difference of antibody function. In practice, we introduce *D*_*E*_, the average Euclidean distance between the embedding of CDR residues of two antibody sequences. The following *in vitro* experiment supports that *D*_*E*_ is related to antigen-binding affinity. We conducted antigen-binding affinity experiment (Enzyme linked immunosorbent assay, ELISA) on eight antibodies mutated from precursor of Pembrolizumab. The results are listed in SI.5. The results show that there is a positive correlation between *D*_*E*_ and OD’s EC50, and there is no significant correlation between *D*_*E*_ and the number of mutations. The experimental results demonstrate that *D*_*E*_ is related to antigen-binding affinity and can be used to guide back mutation for humanized antibodies.

In practice, we perform individual back mutation on each non-CDRs residue for humanized sequence, and then calculate the difference in *D*_*E*_ between the resulting mutated sequence and the original humanized sequence, where *D*_*E*_ is calculated between input sequence and the corresponding parental murine sequence. If Δ*D*_*E*_ is negative, it indicates that the mutation is beneficial for improving antigen-binding affinity. Setting a threshold, perform the corresponding mutation if Δ*D*_*E*_ is less than the threshold. The results are shown in SI.6.a (heavy chain) and SI.6.b (light chain). The number of Avg(iii+iv) decreases from 11.5 to 7.7 when the threshold is set to -2.5, which indicate that back mutation effectively reduces the number of extra mutations.

We further draw a scatter plot of Avg(iii+iv)-i (Figure 4) and observe that the point representing Hu-mAb is above the line connecting two points of Sapiens, while all points representing PLAN are below the line, implying that the number of extra mutations (iii+iv) PLAN is less than other methods when correctly predicting the same number of mutations (i).

**Figure 4.**
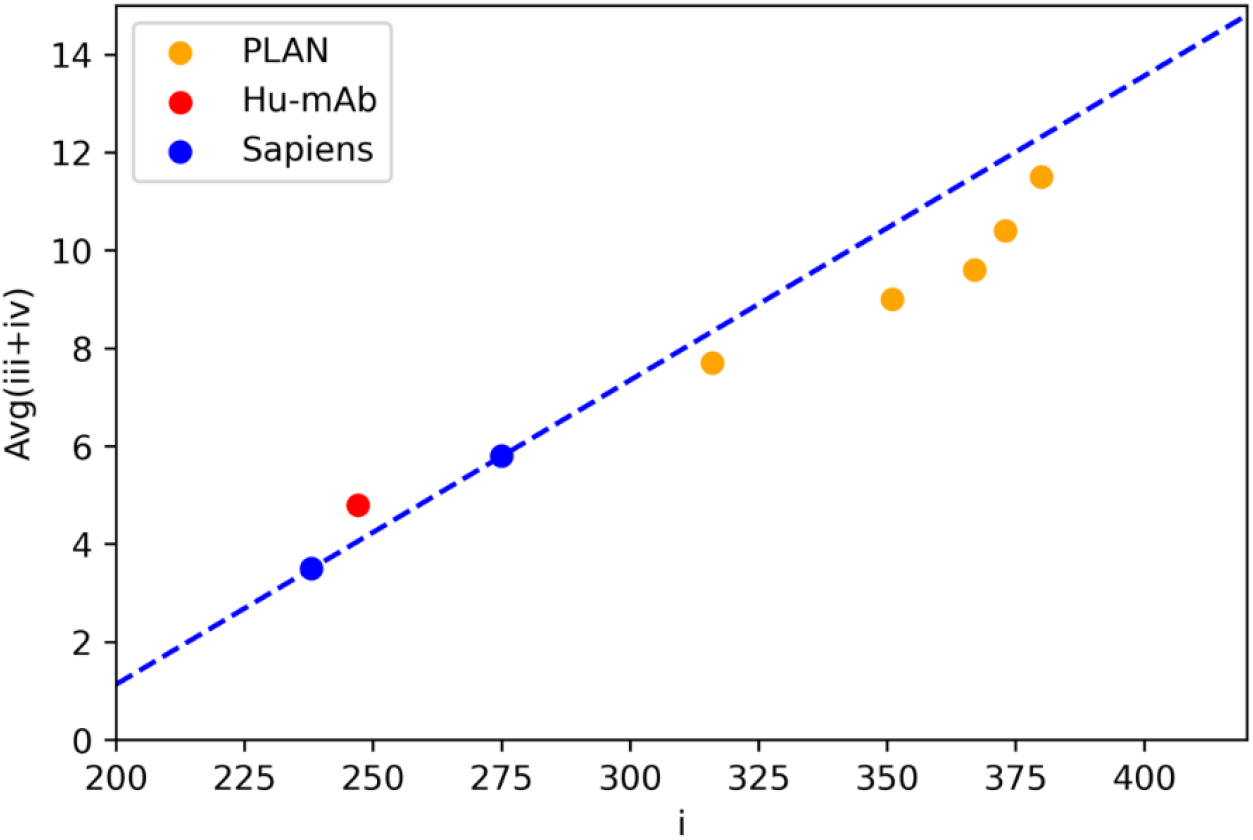
Scatter plot of Avg(iii+iv)-i. This figure depicts a Avg(iii+iv)-i graph and a line connecting two points of Sapiens. It can be observed that the point representing Hu-mAb is above the line, while all points representing PLAN are below the line.

### 3. PLAN-humanized sequences achieve high humanness score

We submit sequences humanized by PLAN, Hu-mAb, and Sapiens to OASis and IgBLAST. The results obtained from OASis and IgBLAST are presented in Table 2 and Table 3. For comparison, we include the results of both experimentally humanized and non-humanized sequences in the table.

**Table 2.**
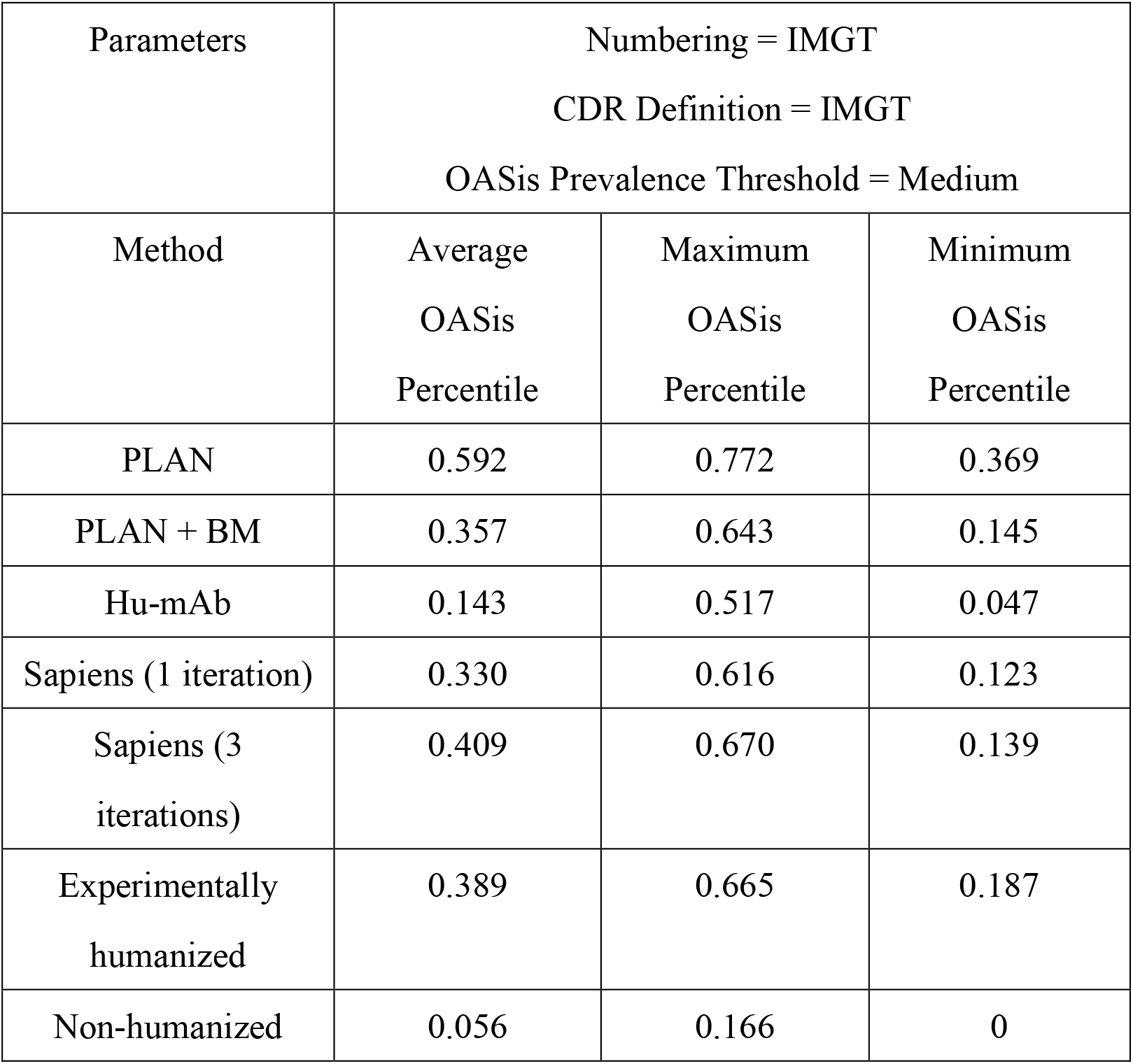
OASis result for each method among 25 antibodies.

**Table 3.**
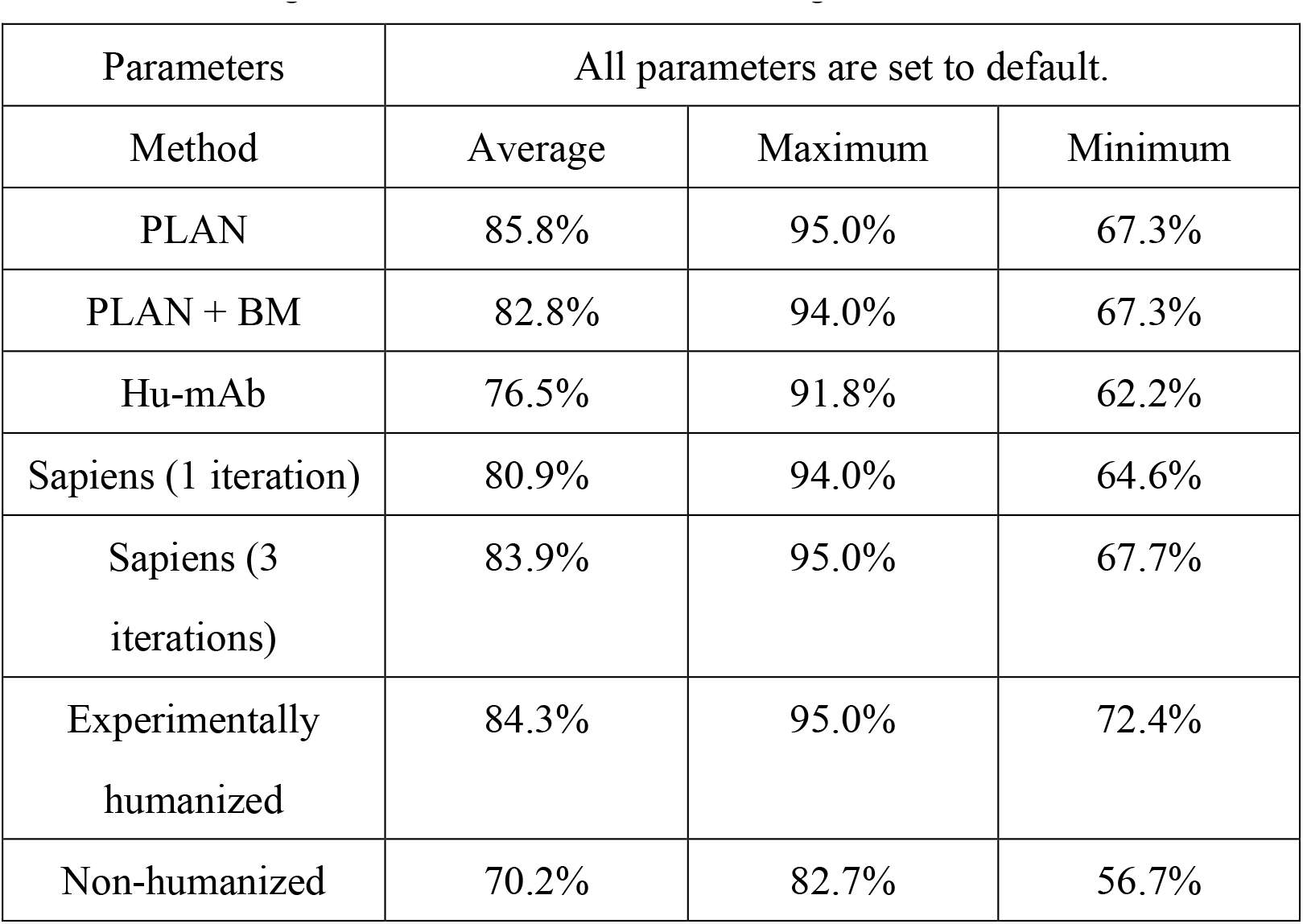
IgBLAST result for each method among 25 antibodies.

| Parameters | All parameters are set to default. |  |  |
| --- | --- | --- | --- |
| Method | Average | Maximum | Minimum |
| PLAN | 85.8% | 95.0% | 67.3% |
| PLAN + BM | 82.8% | 94.0% | 67.3% |
| Hu-mAb | 76.5% | 91.8% | 62.2% |
| Sapiens (1 iteration) | 80.9% | 94.0% | 64.6% |
| Sapiens (3<br>iterations) | 83.9% | 95.0% | 67.7% |
| Experimentally<br>humanized | 84.3% | 95.0% | 72.4% |
| Non-humanized | 70.2% | 82.7% | 56.7% |

According to the results, PLAN-humanized sequences reach the highest average humanness score, even higher than experimentally humanized sequences’ score. The average humanness score of sequences humanized by PLAN+BM is comparable to experimentally humanized sequences’ score, indicating that antibodies processed with back mutation retain acceptable immunogenicity.

### 4. PLAN-humanized sequences achieve comparable stability (according to ΔΔ*G*) to experimentally humanized therapeutics

We run in silico simulation using Flex ddG protocol [30] to compare the performance of PLAN-humanized antibodies to experimentally humanized sequences. We input the experimentally humanized antibody-antigen complex PDB file, and mutate the antibody sequence to the sequence designed by PLAN+BM. For each sequence we run the simulation 3 times under default setting, and we declare comparable binding affinity if the maximum ΔΔ*G* < 1.0 kcal/mol. Out of the 25 experimentally humanized therapeutic sequences, we are able to find 7 antibody-antigen complex PDB file, and 5 of them (Pembrolizumab, Pertuzumab, Eculizumab, Talacoluzumab, Bevacizumab) show comparable binding affinity. The detailed result and its boxplot are shown in Table 4 and Figure 5.

**Table 4.** Flex ddG result for 7 antibodies.

| Antibody | Complex<br>PDB | Maximum<br>$\Delta\Delta G$ | OASis<br>percentile for<br>experimentally<br>humanized<br>antibody | OASis<br>percentile for<br>PLAN-humanized<br>antibody |
| --- | --- | --- | --- | --- |
| Pembrolizumab | 5JXE | 0.1235 | 0.204 | 0.335 |
| Pertuzumab | 1S78 | 0.3266 | 0.322 | 0.322 |
| Certolizumab | 5WUX | 2.1714 | 0.300 | 0.208 |
| Eculizumab | 5I5K | 0.8300 | 0.243 | 0.145 |
| Ligelizumab | 6UQR | 3.1157 | 0.208 | 0.190 |
| Talacoluzumab | 4JZJ | 0.4649 | 0.455 | 0.298 |
| Bevacizumab | 1BJ1 | 0.0200 | 0.328 | 0.166 |

Note: Stabilizing mutations are defined as those with $\Delta\Delta G \leq -1.0$ kcal/mol, neutral as those with $-1.0$ kcal/mol $< \Delta\Delta G < 1.0$ kcal/mol, and destabilizing as those with $\Delta\Delta G \geq 1.0$ kcal/mol [30].

**Figure 5.**
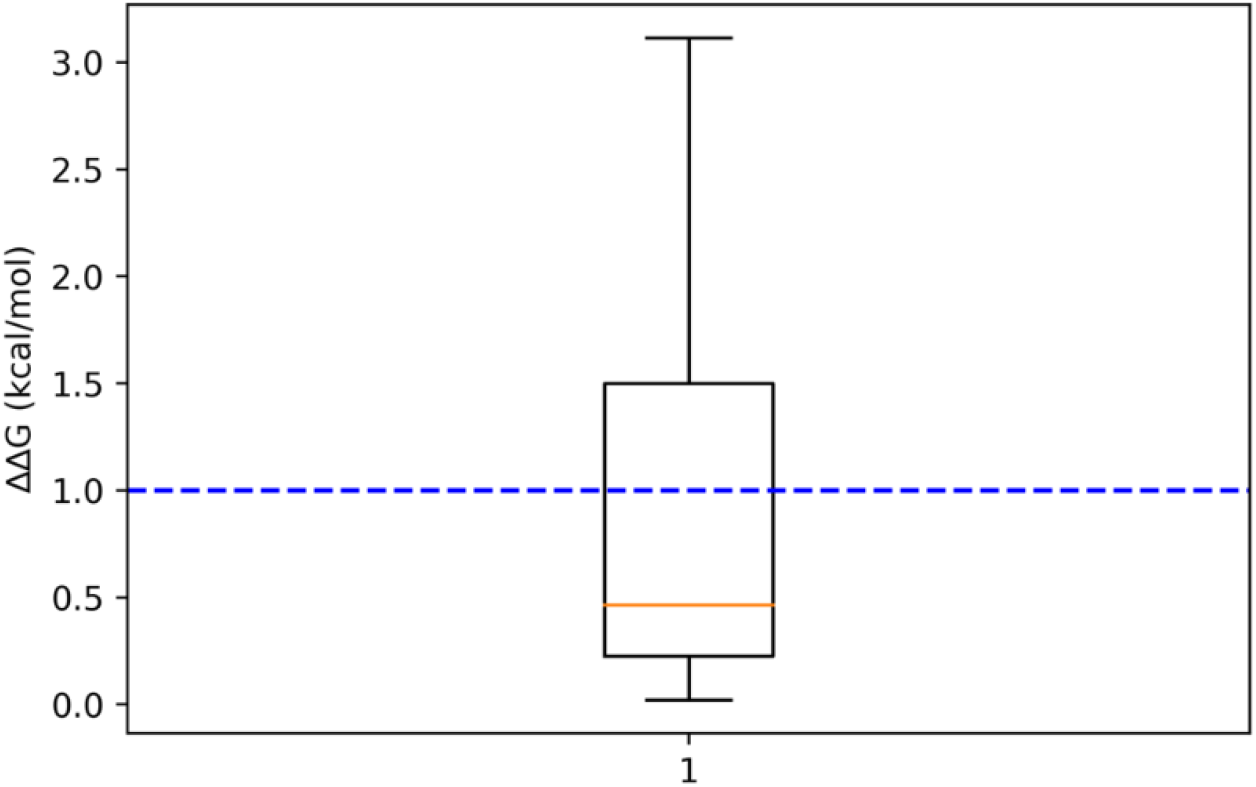
Boxplot of result of Flex ddG for 7 antibodies.

### 5. PLAN achieves high top-3 accuracy rate, enabling efficient mutation scanning on antibody humanization

Assigning candidates to each position can be helpful for subsequent affinity maturation. Since our method sorts all 20 amino acids for each position, we can calculate top 1, top 2, and top 3 accuracy for all positions to verify the validity of the candidates. The result is shown in the Table 5 and Table 6.

**Table 5.**
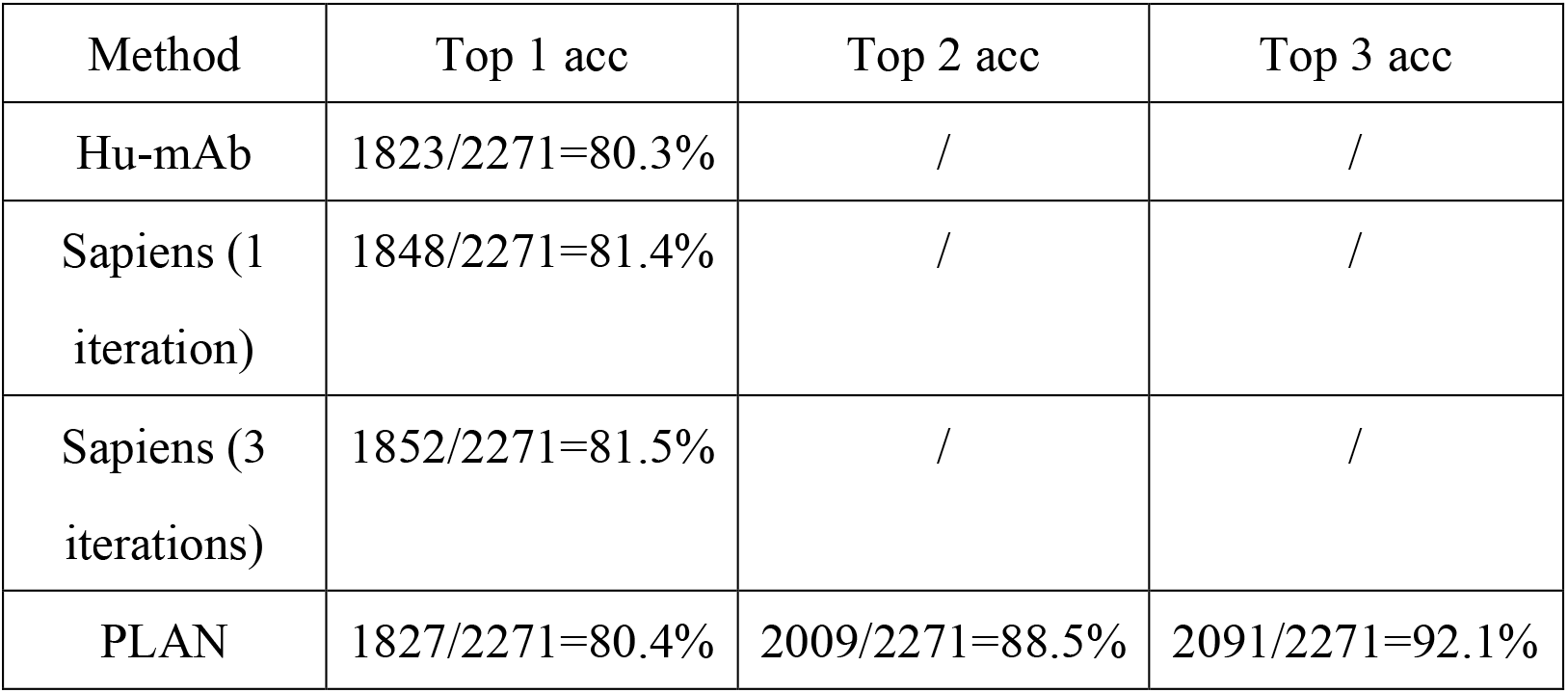
Top 1, top 2, and top 3 accuracy for all positions among 25 antibodies (heavy chain).

**Table 6.**
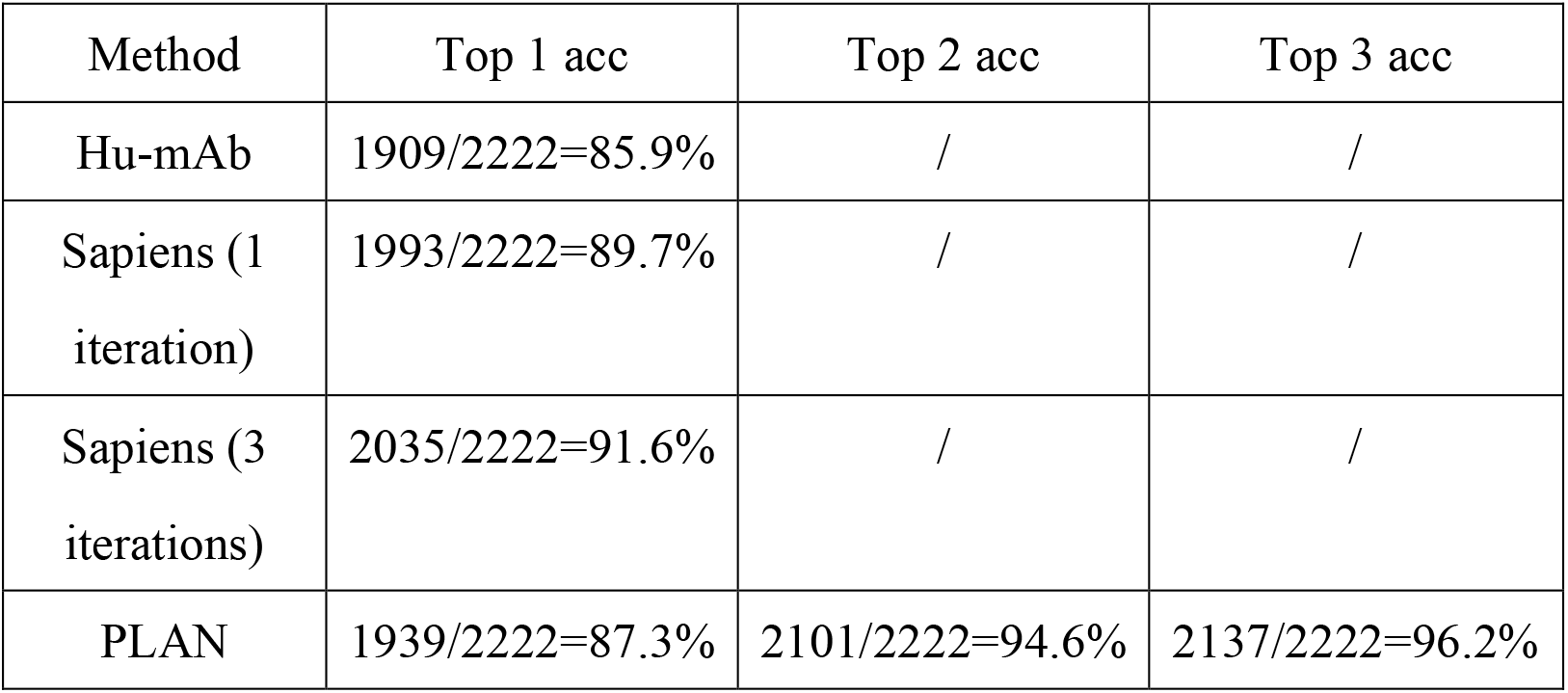
Top 1, top 2, and top 3 accuracy for all positions among 25 antibodies (light chain).

| Method | Top 1 acc | Top 2 acc | Top 3 acc |
| --- | --- | --- | --- |
| Hu-mAb | 1909/2222=85.9% | / | / |
| Sapiens (1 iteration) | 1993/2222=89.7% | / | / |
| Sapiens (3 iterations) | 2035/2222=91.6% | / | / |
| PLAN | 1939/2222=87.3% | 2101/2222=94.6% | 2137/2222=96.2% |

Taking the result of heavy chain as an example, each sequence has 10 positions that fall outside of the top 2 predicted amino acids and has 7 positions that fall outside of the top 3 predicted amino acids on average. It is worth noting that for Lorvotuzumab, an anti-CD56 monoclonal antibody, its top 2 accuracy is 100%. This means that Lorvotuzumab heavy chain can be sampled from top 2 candidates through a well-designed sampling strategy.

Our method can reduce the antibody searching space from approximately 20^200^ (each antibody heavy and light chain framework region has a length of approximately 200, and there are 20 possible choices for each position) to 3^200^, and the reduced searching space contains sequences that have very high sequence identity to experimentally humanized sequences.

## Methods

### 1. Datasets preparation

We prepare our dataset in three steps in this study. First, we randomly select 500,000 human antibody heavy chain sequences and 500,000 human antibody light chain sequences from Observed Antibody Space (OAS) [23], a database contains more than one billion antibody sequences from over 80 different studies (December 2022). Second, to downsize the database while preserving its diversity, we utilize MMseqs2 [28] to cluster the collected antibody sequences and retain only the representative sequence for each cluster. Third, we utilize ESM2 (“esm2_t36_3B_UR50D”) to extract embeddings of the last layer (layer 36) for each of the remaining sequences.

Inspired by germline humanization [29], we also collect all human germline sequences as an optional dataset. Human germline sequences are download from the IMGT database (Jan. 25, 2023). We further select the first one gene for each allele, for the V, D, and J genes. We then combine these genes in an all-against-all fashion and utilize ESM2 (“esm2_t36_3B_UR50D”) to extract embeddings of the last layer (layer 36) for each of the obtained sequences.

### 2. Humanization protocol of PLAN

We suppose that residues adjacent in ESM2’s embedding space have similar biophysical and functional properties. Thus, we humanize the input murine antibody sequence by replacing each residue with another residue appears in a human antibody or germline sequence and is adjacent with the residue in ESM2’s embedding space. In other words, we replace each amino acid in the input sequence with a human-derived version that has similar properties, thereby humanizing it. To make the result more stable, we employ kNN search on K-mer level to implement the above idea.

The following notations are used throughout presentation of this section: *AA*, amino acid; *K-mer*, K-mer; *Seq*, protein sequence; ESM(*AA*)_*seq*_, the embedding generated by ESM2 for amino acid *AA* in protein sequence *Seq*. PLAN humanizes heavy chains and light chains separately, without introducing mutations to the CDR residues. The detailed steps are as follows.

I. Generate embeddings for each of the sequence *Seq* in collected sequence dataset *Dataset* using ESM2 (“esm2_t36_3B_UR50D”, layer 36).

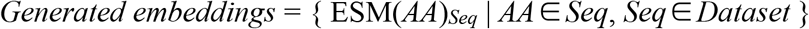
II. Build the humanization kNN pool. Regarding the embedding of each *K-mer* from each sequence in the collected sequence dataset as value and the corresponding *K-mer* as key. The embedding of *K-mer* is defined as the average embedding of each residue within the *K-mer*. Key-value pairs are restored in the kNN pool.

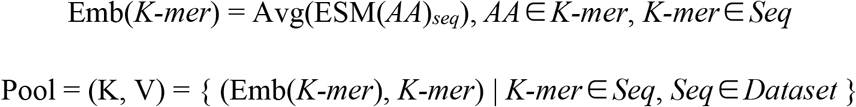
III. Generate embeddings for input murine antibody sequence *Seq*_*m*_ using ESM2 (“esm2_t36_3B_UR50D”, layer 36). For each embedding of *K-mer* in input murine antibody sequence, run kNN search and obtain *k K-mers* from the kNN pool. Count the occurrence of all searched *K-mers* on residue level (e.g., if the first amino acid of a searched *K-mer* is glycine, count one for glycine at the corresponding position in the sequence). Finally, vote for each position, and select the amino acid with the highest count as the result of humanization.

### 3. Back mutation protocol of PLAN

Calculates the Δ*D*_*E*_ for each non-CDRs residue. If Δ*D*_*E*_ is less than a pre-defined threshold, then back mutate it. *D*_*E*_ is defined as the average Euclidean distance between the embedding of CDR residues of a sequence and the embedding of CDR residues of the input murine sequence. *D*_*E*_ measures the similarity of the CDR residues between two protein sequence in ESM2’s embedding space. Based on the previous assumption, the smaller the value of *D*_*E*_, the similar the functional properties of the two protein sequences.

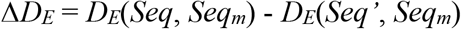

Where *Seq’* back mutate one residue from *Seq*.

### 4. Enzyme linked immunosorbent assay

To evaluate antigen-binding affinity, we conducted enzyme linked immunosorbent assay (ELISA). Antigens were diluted to final concentrations of 2μg/ml, and coated onto 96-well plates and incubated at 4°C for 16h. Samples were washed with 0.05% PBS-T one time and blocked with blocking buffer (PBS containing 5% BSA) at 25°C for 1h. MAbs were diluted in serial dilutions starting at 400 nM with fourth serial dilution, added the plates and incubated at 25°C for 1h. Wells were then incubated with secondary goat anti-human IgG labeled with HRP and TMB substrates. Optical density (OD) was measured by a spectrophotometer at 450nm.

### 5. *In silico* metric of immunogenicity and antigen-binding affinity of antibody

To evaluate the immunogenicity of humanized antibody, we utilize OASis [10] and IgBLAST [27]. OASis separated human and non-human sequences with high accuracy and correlated with clinical immunogenicity; IgBLAST matched the sequence most similar to the query sequence from the human antibody database, whose similarity (sequence identity) to the query sequence indicates the immunogenicity of the query sequence. OASis and IgBLAST take the antibody sequence as input and output a score ranging from 0 to 1 that has been proven to be related to the immunogenicity of the antibody. We compare the humanized sequences obtained from different methods with the experimentally validated humanized sequences by scoring them with OASis (Numbering and CDR Definition scheme = IMGT, OASis prevalence threshold = medium) or IgBLAST (return the highest sequence identity to the queried sequence among matched sequences). If the two scores are close and surpass the score of the parental murine sequence, the immunogenicity of humanized antibody is considered acceptable.

To evaluate the antigen-binding affinity of humanized antibody, we utilize Flex ddg (default settings) [30][35]. The Flex ddG protocol predicts changes in binding free energy upon mutation.

## Discussion

To the best of our knowledge, different from previous work where language models (or generative models) were used to output probabilities of mutated amino acids (or protein sequences) to guide protein optimization, PLAN is the first work to apply BERT-kNN to the field of protein design.

PLAN is leading in two *in silico* humanness scoring. In the early stages of this work, we humanized Pembrolizumab using an early version of PLAN and conducted ELISA to validate antigen-binding affinity *in vitro*. The wet-lab experiment results indicated that while the humanized sequences performed well in *in silico* humanness scoring and retained their ability to bind to antigens, they exhibited a loss of antigen-binding affinity, exceeding a tenfold reduction. We suppose that the reason is that PLAN replaces the amino acids in the parental murine sequence with a certain amino acid that appears in human antibody (or germline) repertoire, but in some cases, there may not be suitable amino acids in the repertoire, which leads to a decrease in antigen-binding affinity after humanization. To solve this problem, we proposed *D*_*E*_ to discover these key residues and back mutate them. The above wet-lab experimental results support the effectiveness of *D*_*E*_ and method of back mutation. Finally, we humanized 7 murine antibodies using PLAN, of which three showed both acceptable immunogenicity and antigen-binding affinity in *in silico* scoring.

In future work, we may ensemble more language models together to obtain stronger residue representation. Structure-based feature extraction methods can also be integrated to obtain more comprehensive features. Structural-based feature extraction methods can also be integrated to obtain more comprehensive amino acid features. Moreover, structural-based feature extraction methods may further help humanization from another dimension, which has chances to alleviate the bias of sequence-based methods. However, it requires structure prediction of antibody sequences that are not in the Protein Data Bank, which suffers bias.

The total top 3 accuracy of PLAN is as high as 95.0%, indicating that there are sequences in the space formed by the top 3 amino acids that are very similar to the sequences verified by wet-lab experiments, and these sequences is worthy for further experimental investigations. To further streamline the *in silico* humanization protocol, we can design fast and effective *in silico* screening methods to identify good sequences from the reduced space. For example, *D*_*E*_ can serve as a *in silico* indicator to guide heuristic search. Molecular dynamics or wet-dry combination methods can also be applied here, as they have high accuracy, but the disadvantage is that their running time is relatively long.

