## Supplementary material for "Antibody Humanization via Protein Language Model and Neighbor Retrieval": Humanization_SI.docx

**Supplementary Information**

Honggang Zou^1^, Rongqing Yuan^1^, Boqiao Lai^2^, Yang Dou^1^, Li Wei^3^, Jinbo Xu^1*^

1 Molecule Mind

2 Toyota Technological Institute at Chicago

3 Shanghai Artificial Intelligence Laboratory

* Corresponding author

1. Individual result for mutation overlapping ratio between experimentally humanized sequences and PLAN-humanized sequences.

| Antibody | Heavy chain | Light chain | Overall |
| --- | --- | --- | --- |
| AntiCD28 | 0.82 | 0.74 | 0.79 |
| Bevacizumab | 0.60 | 0.88 | 0.71 |
| Campath | 0.26 | 0.79 | 0.40 |
| Certolizumab | 0.61 | 0.90 | 0.73 |
| Clazakizumab | 0.63 | 0.91 | 0.76 |
| Crizanlizumab | 0.78 | 0.17 | 0.48 |
| Eculizumab | 0.83 | 0.90 | 0.86 |
| Etaracizumab | 0.69 | 0.56 | 0.61 |
| Herceptin | 0.69 | 0.64 | 0.67 |
| Idarucizumab | 0.33 | 0.50 | 0.38 |
| Ixekizumab | 0.79 | 0.83 | 0.80 |
| Ligelizumab | 0.90 | 0.86 | 0.88 |
| Lorvotuzumab | 0.69 | 0.54 | 0.62 |
| Mogamulizumab | 0.80 | 0.83 | 0.81 |
| Omalizumab | 0.35 | 0.28 | 0.32 |
| Palivizumab | 0.39 | 0.81 | 0.64 |
| Pembrolizumab | 0.65 | 0.10 | 0.40 |
| Pertuzumab | 0.38 | 0.75 | 0.52 |
| Pinatuzumab | 0.39 | 0.04 | 0.25 |
| Refanezumab | 0.71 | 1.00 | 0.85 |
| Reslizumab | 0.95 | 0.85 | 0.90 |
| Rovalpituzuma | 0.57 | 0.54 | 0.55 |
| Solanezumab | 1.00 | 0.60 | 0.85 |
| Talacoluzumab | 0.39 | 0.88 | 0.55 |
| Tocilizumab | 0.70 | 0.89 | 0.79 |

2. Histogram for mutation overlapping ratio between experimentally humanized sequences and PLAN-, Hu-mAb-, and Sapiens-humanized sequences among 25 antibodies.

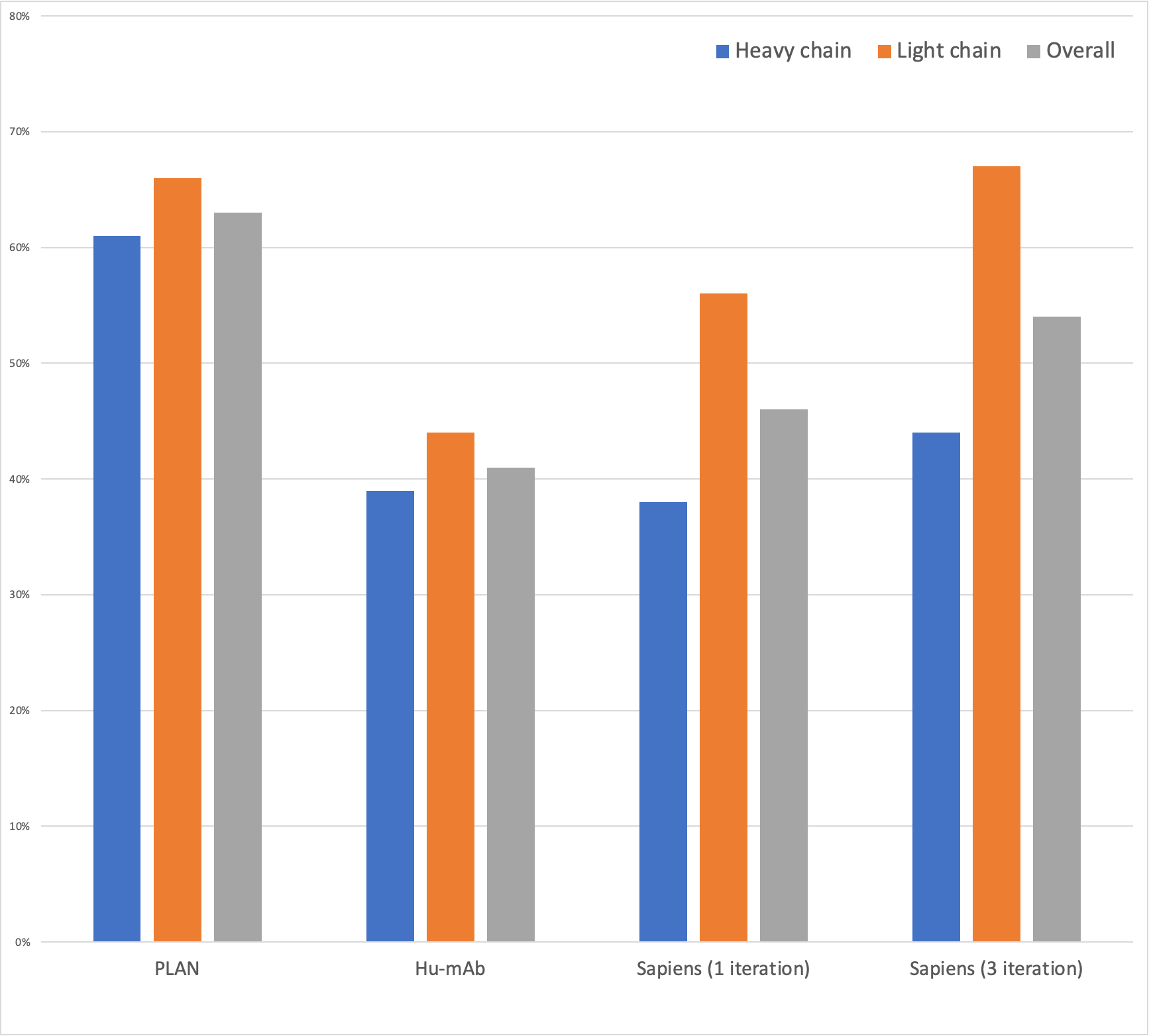

3. An example of classification result for each position.

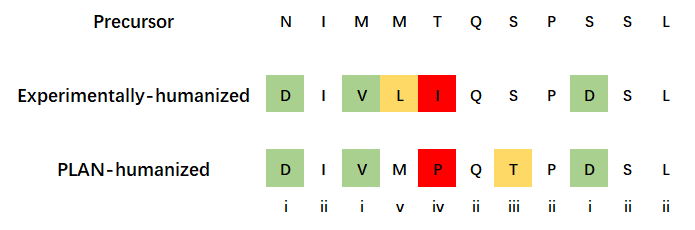

4. Classification result for each position among 25 antibodies.
   a. Classification result for each position among 25 antibodies (heavy chain)

| Method | i ↑ | ii ↑ | iii ↓ | iv ↓ | v ↓ | total |
| --- | --- | --- | --- | --- | --- | --- |
| Hu-mAb | 247 | 1576 | 67 | 53 | 328 | 2271 |
| Sapiens (1 iteration) | 238 | 1610 | 33 | 55 | 335 |  |
| Sapiens (3 iterations) | 275 | 1577 | 66 | 80 | 273 |  |
| PLAN | 380 | 1447 | 196 | 92 | 156 |  |

b. Classification result for each position among 25 antibodies (light chain)

| Method | i ↑ | ii ↑ | iii ↓ | iv ↓ | v ↓ | total |
| --- | --- | --- | --- | --- | --- | --- |
| Hu-mAb | 206 | 1703 | 50 | 10 | 253 | 2222 |
| Sapiens (1 iteration) | 264 | 1729 | 24 | 30 | 175 |  |
| Sapiens (3 iterations) | 312 | 1723 | 30 | 34 | 123 |  |
| PLAN | 307 | 1632 | 121 | 48 | 114 |  |

5. Result of EC50, *D_E_*, and mutation number on eight antibodies mutated from precursor of Pembrolizumab and Pembrolizumab itself.

| Antibody | EC50(nM) | *D_E_* | Mutation number |
| --- | --- | --- | --- |
| mPembrolizumab_0 | ∞ | 114.5572 | 42 |
| mPembrolizumab_1 | ∞ | 114.7343 | 43 |
| mPembrolizumab_2 | ∞ | 114.9969 | 43 |
| mPembrolizumab_3 | ∞ | 116.9213 | 43 |
| mPembrolizumab_4 | 568 | 112.1118 | 43 |
| mPembrolizumab_5 | 773 | 113.6161 | 44 |
| mPembrolizumab_6 | 97 | 111.5790 | 42 |
| mPembrolizumab_7 | 46 | 106.0776 | 41 |
| Pembrolizumab | 0.97 | 102.4652 | 44 |

6. Classification result after *D_E_* guided back mutation for each position among 25 antibodies.
   a. Classification result after *D_E_* guided back mutation for each position among 25 antibodies (heavy chain).

| Method | i ↑ | ii ↑ | iii ↓ | iv ↓ | v ↓ | Avg (iii + iv) ↓ | total |
| --- | --- | --- | --- | --- | --- | --- | --- |
| PLAN | 380 | 1447 | 196 | 92 | 156 | 11.5 | 2271 |
| PLAN + BM (threshold = -4) | 373 | 1474 | 169 | 90 | 165 | 10.4 |  |
| PLAN + BM (threshold = -3.5) | 367 | 1490 | 153 | 87 | 174 | 9.6 |  |
| PLAN + BM (threshold = -3) | 351 | 1503 | 140 | 84 | 193 | 9.0 |  |
| PLAN + BM (threshold = -2.5) | 316 | 1527 | 116 | 76 | 236 | 7.7 |  |
| Hu-mAb | 247 | 1576 | 67 | 53 | 328 | 4.8 |  |
| Sapiens (1 iteration) | 238 | 1610 | 33 | 55 | 335 | 3.5 |  |
| Sapiens (3 iterations) | 275 | 1577 | 66 | 80 | 273 | 5.8 |  |

b. Classification result after *D_E_* guided back mutation for each position among 25 antibodies (light chain).

| Method | i ↑ | ii ↑ | iii ↓ | iv ↓ | v ↓ | Avg (iii + iv) ↓ | total |
| --- | --- | --- | --- | --- | --- | --- | --- |
| PLAN | 307 | 1632 | 121 | 48 | 114 | 6.8 | 2222 |
| PLAN + BM (threshold = -3.25) | 283 | 1655 | 98 | 45 | 141 | 5.7 |  |
| PLAN + BM (threshold = -3) | 275 | 1659 | 94 | 43 | 151 | 5.5 |  |
| PLAN + BM (threshold = -2.75) | 261 | 1674 | 79 | 42 | 166 | 4.8 |  |
| PLAN + BM (threshold = -2.5) | 245 | 1687 | 66 | 39 | 185 | 4.2 |  |
| Hu-mAb | 206 | 1703 | 50 | 10 | 253 | 2.4 |  |
| Sapiens (1 iteration) | 264 | 1729 | 24 | 30 | 175 | 2.2 |  |
| Sapiens (3 iterations) | 312 | 1723 | 30 | 34 | 123 | 2.6 |  |

Where Avg (iii + iv) denotes the number of (iii + iv) per sequence, representing number of extra mutations per sequence.
